# Identification of Gene Regulation Models from Single-Cell Data

**DOI:** 10.1101/231415

**Authors:** Lisa Weber, William Raymond, Brian Munsky

**Affiliations:** Department of Chemical and Biological Engineering, Colorado State University, Fort Collins, CO; School of Biomedical Engineering, Colorado State University, Fort Collins, CO

**Keywords:** gene expression models, stochastic gene expression, single-cell data, model inference

## Abstract

In quantitative analyses of biological processes, one may use many different scales of models (e.g., spatial or non-spatial, deterministic or stochastic, time-varying or at steady-state) or many different approaches to match models to experimental data (e.g., model fitting or parameter uncertainty/sloppiness quantification with different experiment designs). These different analyses can lead to surprisingly different results, even when applied to the same data and the same model. We use a simplified gene regulation model to illustrate many of these concerns, especially for ODE analyses of deterministic processes, chemical master equation and finite state projection analyses of heterogeneous processes, and stochastic simulations. For each analysis, we employ Matlab and Python software to consider a time-dependent input signal (e.g., a kinase nuclear translocation) and several model hypotheses, along with simulated single-cell data. We illustrate different approaches (e.g., deterministic and stochastic) to identify the mechanisms and parameters of the same model from the same simulated data. For each approach, we explore how uncertainty in parameter space varies with respect to the chosen analysis approach or specific experiment design. We conclude with a discussion of how our simulated results relate to the integration of experimental and computational investigations to explore signal-activated gene expression models in yeast [1] and human cells [2]‡.

PACS numbers: 87.10.+e, 87.15.Aa, 05.10.Gg, 05.40.Ca,02.50.-r

Submitted to: Phys. Biol.

## 1. Introduction

Recent years have led to rapid advances in the capabilities to measure gene regulatory phenomena at the level of single-cells and in fluctuating conditions. These techniques include image-based strategies, where fluorescence markers are used to highlight specific DNA, RNA, or protein molecules of interest [4, 5]; cytometry-based approaches, where cellular properties, especially size or fluorescence, are rapidly measured one cell at a time [6]; mass-spectrometry based approaches, where cellular proteins are digested into peptide fragments whose distinct mass or charge spectra can be analyzed quantitatively [7]; and sequencing approaches, where the abundances of specific RNA or DNA molecules are quantified on a single-cell basis [8]. In this article, we focus on the quantitative analysis of one specific technique, known as single-molecule Fluorescence *in situ* Hybridization (smFISH) [5, 9], which provides precise and reproducible measurements of the number and locations of individual RNA molecules within single cells. In the technique of smFISH, researchers tile individual molecules of endogenous RNA with many complementary DNA oligomers. Each RNA-binding oligomer contains one or more fluorescent dyes, such that the entire multi-labeled RNA molecule appears as a bright diffraction limited spot under the fluorescence microscope. Counting the integer numbers of these spots for many single cells in large populations at many times and in the same conditions, one can capture the temporal distributions of gene regulation activity for those conditions. By using microfluidics or by adding or removing stimuli, one can observe how single-cell statistics are affected by transitions from one environmental condition to another [1, 10, 11].

Utilizing experiments such as smFISH, many researchers have quantified time-varying and bursty gene expression at the single-cell level, and many models have been developed to capture these dynamics (e.g., see recent reviews at [12, 13, 14]). In this article, we define a small class of single-cell models which have previously been shown to capture and predict smFISH data [1, 10, 11], and we employ several computational analyses in attempts to identify these models and their parameters from various sets of simulated data.

### 1.1 A Standard Computational Toolbox for Quantitative Biology

In the following paragraphs, we introduce a handful of computational tools that are frequently used in the analysis and fitting of quantitative gene regulation models.

#### Ordinary Differential Equations

One of the most common approaches to describe the dynamic behavior of gene regulation is to use coupled sets of Ordinary Differential Equations (ODEs). For example, the ODE related to mRNA transcription is a mathematical equation which describes the rate of change in the number of mRNA over time, taking into consideration both production and degradation of mRNA molecules. For most biological processes of interest, analytical solutions to the corresponding ODEs may not be possible and numerical techniques may be necessary. In this article, we will deal with reaction rates that change over time, for which numerical integration is required.

#### Parameter Fitting and the Metropolis-Hastings Algorithm

A common task in quantitative biology is to fit parameters of a model to match experimental data. Due to the stochastic nature of biological systems, in addition to the fact that experimental data is noisy and limited in quantity, exact estimation of parameters is impossible. Having a quantitative understanding of parameter uncertainties in light of existing data is crucial for predictive modeling. To accomplish this goal, Markov Chain Monte Carlo (MCMC) analyses, such as the Metropolis-Hastings (MH) algorithm [15], are frequently used to provide unbiased samples of parameter space and to estimate parameter uncertainties. In particular, the MH algorithm samples parameter space by adding random perturbations to parameters and then calculating the likelihood of the data, given the new parameter set. The new parameter set is then discarded unless the likelihood exceeds an appropriately chosen (albeit) random acceptance threshold. Like any parameter fitting routine, the MH algorithm has several tunable parameters. The *MH chain* is the sequence of parameter sets that have been accepted during the random walk through parameter space. The *proposal distribution* is a probabilistic rule by which a new parameter set is generated from the old. The *burn-in* period is the number of MH samples that are ignored to allow the MH chain to relax away from the (potentially poor) initial parameter guess. The *chain length* is the total number of MH samples that are recorded following the burn-in period. The *thin rate* is the number of accepted samples that are ignored between those that are recorded (i.e., every 10^th^ sample). See [3] for a more in depth discussion of Monte Carlo analyses (Chapters 10 and 16), parameter estimation (Chapter 13), and sensitivity analyses (Chapter 14) as applied to models in quantitative biology.

#### The Stochastic Simulation Algorithm

Unlike most dynamical problems in science and engineering, gene regulation is inherently subject to discrete stochastic behavior arising from the single-molecule nature of genes within individual cells. The standard approach to analyze such discrete stochastic processes is to apply the kinetic Monte Carlo algorithm known as (Gillespie’s) *Stochastic Simulation Algorithm* (SSA) [16, 17], which uses a combination of stochastic reaction rates (i.e., propensity functions) and reaction rules (i.e., stoichiometry vectors) to simulate the stochastic trajectories. This algorithm updates the state of the system after each random reaction and assumes exponentially-distributed waiting times between every event. In this article, we apply the SSA to simulate many possible stochastic trajectories for the switching of gene activity states coupled with mRNA production and degradation.

#### The Finite State Projection Method

In addition to using the SSA to generate stochastic trajectories for the gene regulation processes, we also explore direct computations of the underlying probability distributions for these processes. These distributions evolve according to the infinite dimensional linear equation known as the Chemical Master Equation (CME) [18, 19]. Because the CME has no exact solution for most models, we employ the finite state projection (FSP) method to approximate the CME solution and to estimate the full joint probability distributions for gene switching behavior and transcript accumulation [20]. Several recent reviews of the FSP approach and its applications can be found at [14, 21, 22, 23]. A key advantage of the CME and FSP formulation is that they provide systematic approaches to compute and compare the likelihood of data arising from multiple models [1, 10, 21]. With this in mind, we use the Metropolis-Hastings algorithm in conjunction with the FSP analysis to attain more precise constraints on the parameters when fit to simulated data [11].

#### Experiment Design

The successful identification of a model from data requires not just good data and appropriate modeling tools, but the experiment itself also needs to be conducted using a well-designed and informative condition [23, 24, 25, 26, 27, 28, 29]. With this in mind, we will explore the importance of experiment design, and we will show that different experiments (e.g., different input signals and different measurement times) and different analyses (e.g., ODEs or the FSP) yield different results for the constraint of system parameters.

In the following section, we introduce a specific gene regulation model and its underlying assumptions. Then, in Section 3, we discuss the simulated data from which we seek to identify our model parameters. In Section 4, we demonstrate the implementation of the above tools to this model and different sets of simulated data. Finally, in Section 5, we provide a brief summary of our results and conclusions, and we introduce a user-friendly Python-based toolbox with a graphical-user-interface to allow readers to reproduce all of the described analyses. All analyses discussed in this article have been implemented in Matlab and Python codes, and the codes are available for download at: https://github.com/MunskyGroup/WeberPB

## 2. Gene Regulation Model Description

We begin by specifying a simple semi-mechanistic model for gene regulation. Figure 1 illustrates this model, which is a generalization of the standard bursting gene expression model discussed in [12, 13, 30]. The model consists of a single gene with three possible states {*S*_1_, *S*_2_, and *S*_3_}, each of which represents a different configuration of transcription factor binding or chromatin conformation. Similar three-state models have been used previously in the context of time-varying inputs to capture and predict signal-activated single-cell gene expression in yeast [1, 11] and human [10] cells. The model assumptions are as described with additional detail in Subsection 2.1.

**Figure 1.**
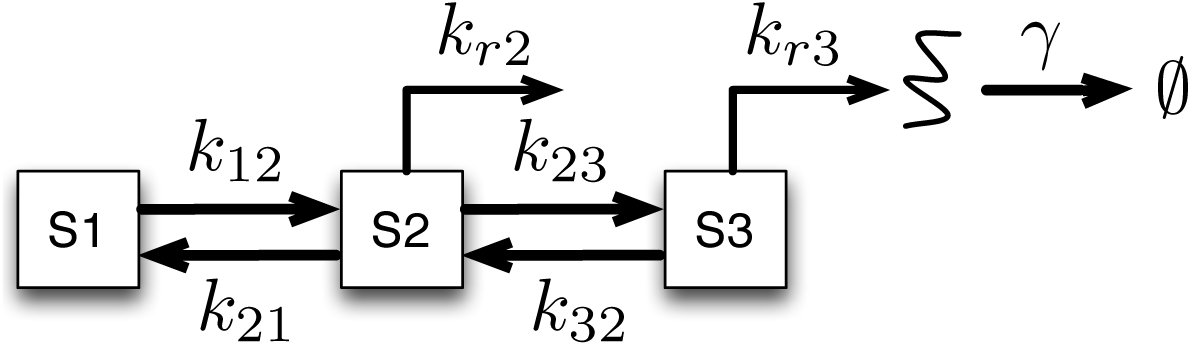
Schematic of a three-state gene regulation model. State 1 is transcriptionally inactive, whereas states 2 and 3 are transcriptionally active with transcription rates *k*_*r*2_ and *k*_*r*3_, respectively. Parameters *k*_*ij*_ denote the (potentially time-varying) transition rates between states *S*_*i*_ and *S*_*j*_.

### 2.1 Assumptions of the Three-State Bursting Gene Expression Model

First, we define the conformational states {*S*_1_, *S*_2_, and *S*_3_} of the gene and the production and degradation of mRNA. In this model, *S*_1_ is a transcriptionally inactive (‘OFF’) state, whereas *S*_2_ and *S*_3_ are transcriptionally active (‘ON’) states. The transcription rates for *S*_2_ and *S*_3_ are *k*_*r*2_ and *k*_*r*3_, respectively. Degradation of mRNA species, *R*, is assumed to be a first order process with rate *γ*.

Next, we define the transition dynamics that allow the gene to switch among these states. In the three-state model, there are four state transition events: *S*_*i*_ *→ S*_*j*_, each with a corresponding transition rate of *k*_*ij*_(*t*). Three of the four transition rates are assumed to be constant with respect to time and the activating kinase: *k*_*ij*_(*t*) = *k*_*ij*_(0). The gene is activated by a time-varying kinase, *Y*_*i*_(*t*), which acts as an input to the system. In practice, such signals can be controlled experimentally through modulation of external stimuli [1, 10, 11, 31, 32, 33]. For illustrative purposes, we initially assume a sinusoidal input signal given by:

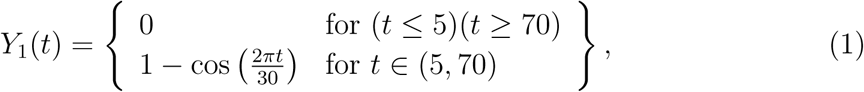

where the input signal (in arbitrary units of concentration) begins at time *t* = 5 min and ends at time *t* = 70 min. We assume exactly one of the four transition rates is linearly dependent upon the activating kinase as follows [1]:

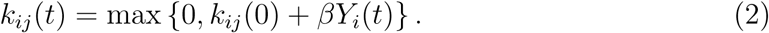

For the combination *i* < *j* (i.e., when the reaction leads to a state with a higher index), the value of *k*_*ij*_ (0) is negative and *β* is positive. Conversely, when *i > j* (i.e., when the reaction leads to a state with a lower index), the value of *k*_*ij*_ (0) is positive and *β* is negative. This choice gives rise to a thresholded activation response in which *k*_*ij*_ (*t*) is zero until the kinase exceeds or drops below a specific level. This definition leads to activation, either through direct activation when *i* < *j* or indirect loss of repression when *i* > *j*.

One of our goals in this study is to determine if it is possible to identify the mechanism of action, i.e., to determine which *k*_*ij*_ depends upon the time-varying kinase signal. We define the model number, *N*_*m*_, as the integer that specifies which mechanism depends upon the activating kinase. The model numbers and the corresponding input-dependent transition rates are as follows:

Model 1: *k*_12_(*t*); Model 2: *k*_23_(*t*); Model 3: *k*_21_(*t*); Model 4: *k*_32_(*t*).

Now that we have specified the assumptions needed to define the gene states, mRNA levels, and the associated dynamics, we can write a set of four ODEs to describe this model:

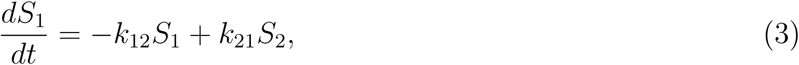

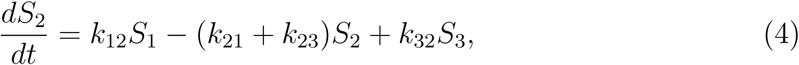

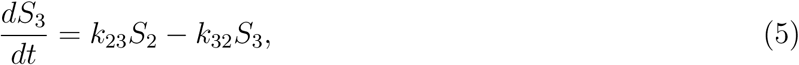

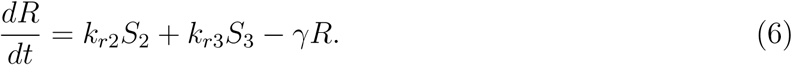

These equations are easily integrated in MATLAB using the stiff ODE solver, ode15s.

Finally, for parameter estimation, we assume that all rates have an independent prior given by a lognormal distribution with a logarithmic mean of 𝔼{log_10_(*λ*)} = 0 and a wide logarithmic variance of 𝔼{(log_10_(*λ*))^2^} = 4. Having defined the reactions and priors on all parameters, in the next section we will generate some simulated data and attempt to fit our model, after which we will explore uncertainty in parameter space, and make model predictions in light of these datasets.

## 3. Single-cell Data Collection

The technique of smFISH [5, 9] makes it possible to measure the number of mRNA molecules within hundreds or thousands of individual cells at specific instances in time during a dynamic response from one condition to another. For example, in [1, 11], the authors measured and found a quantitative FSP-based model to predict the statistics of *STL1, CTT1*, and *HSP12* mRNA expression in yeast cells every two to five minutes following osmotic shock and temporal activation of the HOG1p kinase. Similarly, in [10], the authors measured time-varying p-ERK kinase and its effect on *c-Fos* expression in human-derived U20S cells.

To generate simulated data of this form, we assume a “true” model, which corresponds to Model 2, and for which the parameters are presented in vector form as:

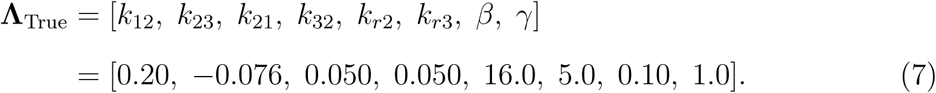

Throughout this article, all rate parameters will be assumed to have units of (minutes)^−1^, and *β* has units of (minutes)^−1^(concentration)^−1^.

At the initial time (*t* = 0), we assume that all cells are in state *S*_1_ and that there are no mRNA (*R* = 0), meaning that all cells are in a transcriptionally inactive state. This initial condition was chosen arbitrarily, but matches to observed dynamics for many genes (e.g., *STL1, CTT1, HSP12, c-Fos*), which are transcriptionally silent in the absence of their activating kinase signal [1, 10]. We use the FSP approach [14, 21, 22, 23, 34] with the true model and known initial condition to compute the time-varying mRNA probability distribution at each combination of time and input condition. We then use the FSP solution to generate histograms of 100 exact independent random samples (i.e., one data point for each simulated cell) at ten equally spaced times between zero and 100 minutes. In the following sections, we will use various computational approaches to infer model parameters from these time-varying histograms.

## 4. Results

The goal of our study is to determine if simulated data is sufficient to identify the model and its parameters, and if the identified parameters can be used to predict the system’s response under new experimental conditions.

### 4.1. Simulation of ODE Models

We first use a deterministic approach and solve the simple system of ODEs listed in Eqs. 3 −6 to capture the bulk gene regulation dynamics. For a given model and parameter set, these ODEs generate the mean gene expression as a function of time. Figure 2 shows the predicted mean gene expression as a function of time for Model 2 (i.e., where *k*_23_ is time-dependent), assuming the input signal in Eqn. 1, and two different arbitrary parameter sets (blue and magenta) as given in the caption.

**Figure 2.**
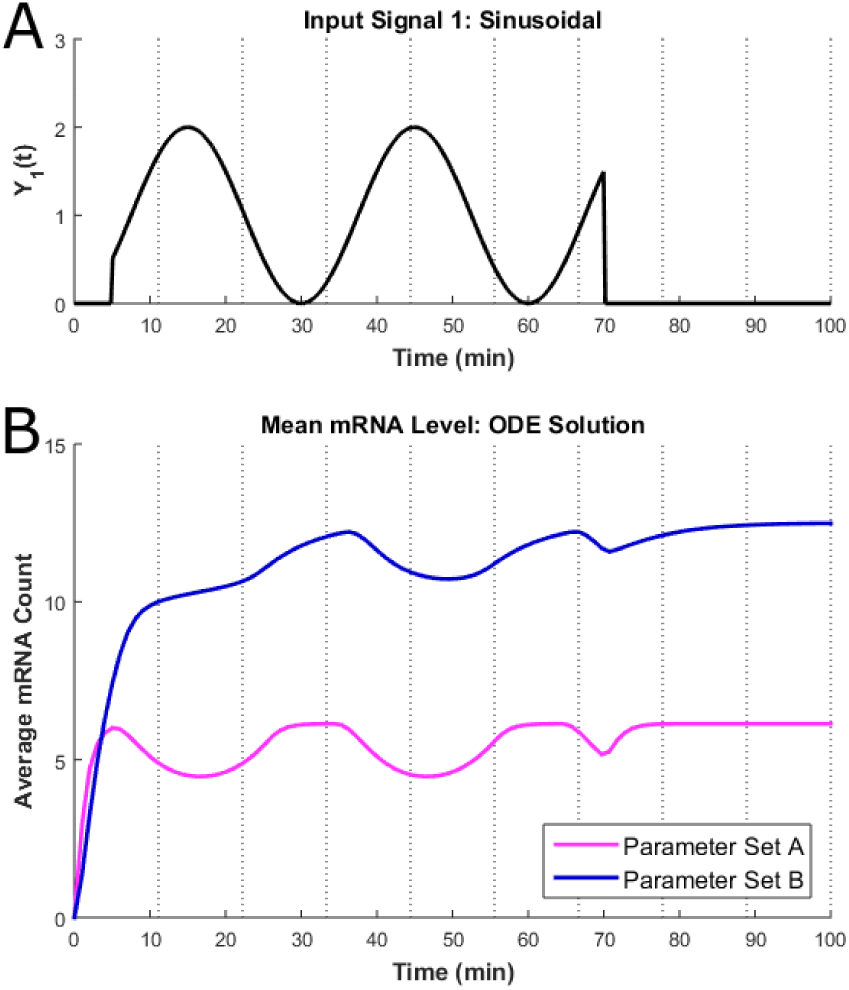
(A) Input signal, *Y*_1_(*t*) in arbitrary units of concentration (AUC) versus time, as defined in Eqn. 1. (B) Deterministic ODE solution for the mean gene expression as a function of time for the three-state gene regulation model shown in Fig. 1, as outlined in Section 4.1. Analyses are shown for Model 2 (*k*_23_ is time dependent), and the two illustrated parameter sets are **Λ**_**A**_ = [3, −0.4, 2.5, 1, 9, 2, 1, 0.8] and **Λ**_**B**_ = [1, −0.6, 1, 0.2, 5, 2, 1, 0.2].

Examining how different parameters affect system behavior reveals qualitative insight about how rate parameters control the underlying process dynamics. For example, Fig. 2 illustrates that faster rates for transcription and degradation (*k*_*r*2_ and *γ*) enable gene expression to track the input signal more closely (Fig. 2, magenta line), whereas slower rates result in greater averaging and lag behind the input signal (Fig. 2, blue line). Biologically, evolution could tune the rates of these reactions to allow cells to respond more quickly to important fluctuations (e.g., sustained stresses) while rejecting faster environmental disturbances (e.g., short-lived transient fluctuations). In addition, such dependence of response dynamics on parameters makes it feasible to perform an inverse analysis (i.e., to estimate parameters from quantitative observations of response dynamics), as discussed in the following subsection.

### 4.2. Maximum Likelihood Estimation for ODE Models

Next, we use the ODE model defined in Eqns. 3-6 and search parameter space to find a parameter set that maximizes the likelihood of the simulated dataset. As described in Section 3, the data were generated at ten equally-spaced time points between zero and 100 minutes, assuming the known input signal, *Y*_1_(*t*), as given in Eqn. 1. Prior to fitting with the ODE model, we computed the mean, standard deviation, and standard error of the mean (see Fig. 3).

**Figure 3.**
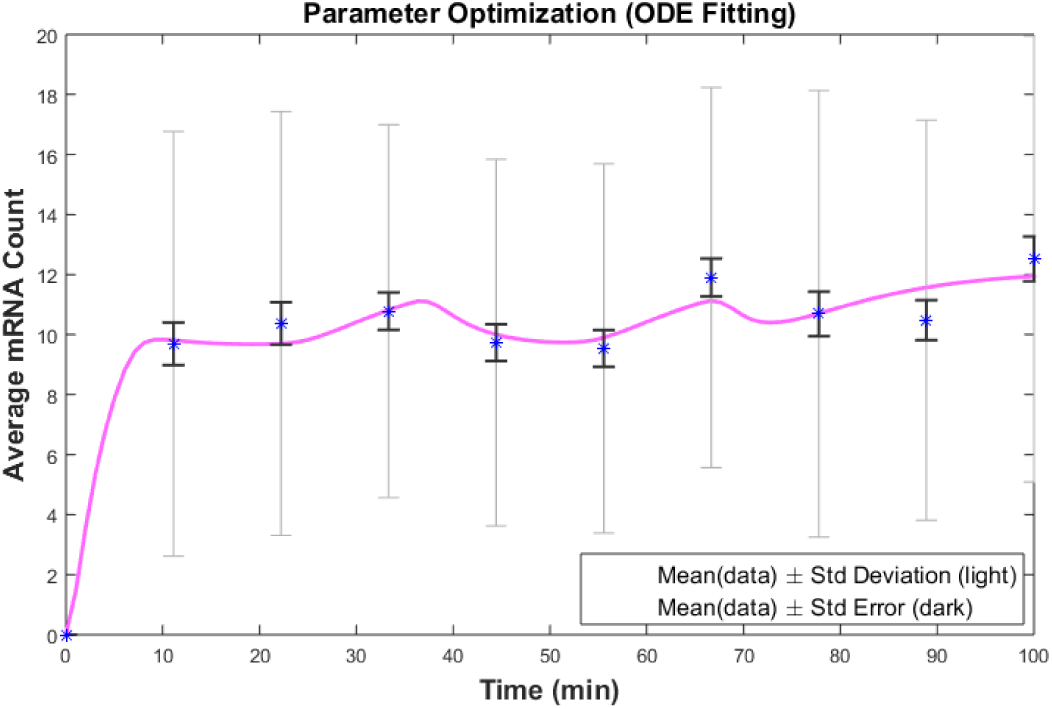
Mean gene expression as a function of time for Model 2 after performing optimization to determine the parameter set with the best fit to the averaged data. The light error bars represent the mean of the data ± one standard deviation. The dark error bars represent the mean of the data *±* the standard error (100 independent samples per time point). The best ODE-fit parameter set for this model is given in Eqn. 14.

Before we can fit the model to the data, we must first compute the likelihood of the data, given the model’s parameters. Consider a set of experimental data, including *N*_*i*_ cells at each time *t*_*i*_. Let 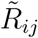 denote the number of mRNA in the *j*^th^ cell at the *i*^th^ time point. The measured sample mean at each time is given by:

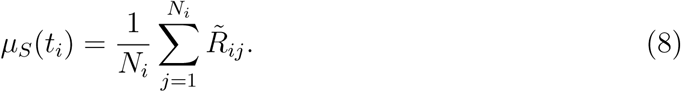

To fit the ODE model to the measured sample mean, one must quantify how well one would expect the sample mean to match to the predictions made by the ODE analysis. This question is easily answered by invoking the central limit theorem (CLT) [35], which applies provided that the true distribution has finite variance. According to the CLT, when many independent and identically distributed measurements are taken from the same distribution, the average of those measurements will tend to a Gaussian random variable with mean equal to the true mean and variance equal to the true variance divided by the number of samples (or squared standard error). In turn, the squared standard error can be estimated from the measured histograms according to:

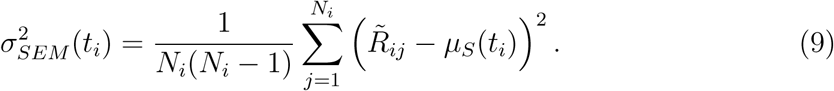

Provided that the underlying distribution has a finite true variance and if enough sample measurements are collected, the likelihood to observe the measured sample mean can be estimated using the normal distribution:

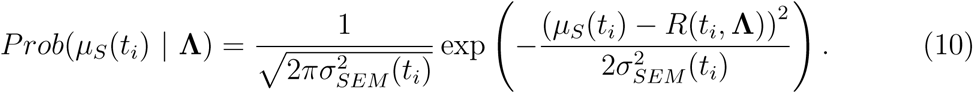

Our assumed lognormal prior distribution (𝔼{*log*_10_(*λ*)} = 0, 𝔼{(*log*_10_(*λ*))^2^} = 4) for each parameter can be written as:

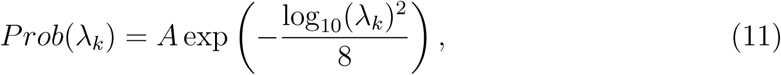

where *A* is a normalization constant. The probability of a parameter set **Λ**, given the data, can be written according to Bayes’ rule, (i.e., *P* (**Λ***|***D**) = *P* (**D***|***Λ**)*P* (**Λ**)*/P* (**D**), where **Λ** denotes the model and **D** denotes the data) as:

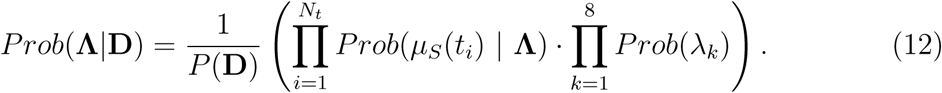

Here, *N*_*t*_ is the number of time points, and *P* (**D**) is a normalization constant. Taking the logarithm of *Prob*(**Λ***|***D**) and substituting in the expressions for *Prob*(*µ*_*S*_(*t*_*i*_) *|* **Λ**) and *Prob*(**Λ***|***D**) above yields:

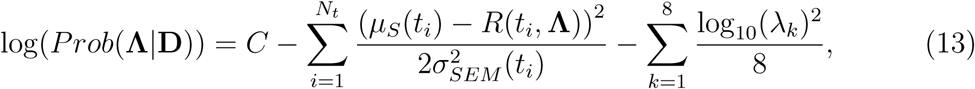

where the first summation is the chi-squared function, which quantifies the likelihood of the data given the model, and the second summation relates to the prior probability to select those parameters in the absence of any data. The variable *C* is a normalization constant that does not depend upon any of the model parameters, and it can be ignored for the purpose of model comparison or likelihood maximization. For small data sets, the second term will dominate, but as more data is collected, the first summation will take on greater importance.

We fit all four possible models (each with a different input-dependent transition rate) by maximizing the likelihood of the model given the data, as specified in Eqn. 13. For this, we use standard optimization routines, including Matlab’s *fminsearch*, which attempts to find the unconstrained minimum of the objective function using a Nelder-Mead simplex algorithm [36, 37]. We have also utilized a customized simulated annealing algorithm [38]. Knowing the signs of all parameters, we converted them to positive values, and we conducted all searches in logarithmic space. This choice is advantageous because (i) it allows us to improve efficiency by removing need to check and satisfy parameter positivity constraints, and (ii) it allows for parameter step magnitudes to be relative to their current values, even when relative magnitudes of parameters are unknown at the start of the fitting process. To increase our chances of finding a global maximum of the likelihood function (and to avoid getting stuck in local maxima associated with particular parameter guesses), we restarted all searches using thirty different random initial guesses per model. We selected the local maximum that provide the greatest likelihood of the model, given the data. Figure 3 shows the result for such a fit using the “correct” model (i.e., Model 2) in which the rate *k*_23_ was assumed to be time-dependent. The corresponding best-fit parameter set for Model 2 using the ODE fit was:

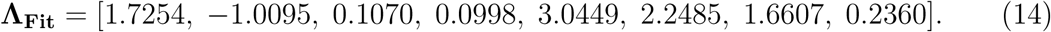

These parameters are very different from the “true” parameters given in Eqn. 7, suggesting that although the ODE analysis provides excellent fits, it is insufficient for identification of the parameters.

### 4.3. Parameter Uncertainty Estimation for ODE Models

When maximizing the ODE-based likelihood of the model, given the data, we saw that it was possible to find many combinations of models and parameters that match exceptionally well to the simulated data. For example, Models 2 and 4 fit equally well to the data generated from Model 2, meaning that the ODE analysis was insufficient for model discrimination. We next explored the uncertainty in parameter space in order to understand how well the averaged experimental data was able to constrain the parameters of the chosen model. To do this, we implemented the Metropolis-Hastings (MH) algorithm (see Section 1.1) to explore posterior parameter uncertainty in the model, given the simulated experimental data and the likelihood function in Eqn. 13.

For our implementation of the MH search, we chose a proposal function in which we select each parameter with probability of 0.5, and, for those selected, we change their logarithmic value by adding a Gaussian-distributed perturbation with a mean of zero and a standard deviation of 0.2. In Matlab, this can be scripted as:

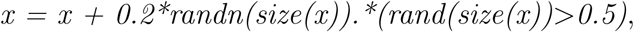

where ‘x’ is the vector of parameters in logarithmic space. For the MH search of parameter space, we used a total chain length of 750,000 parameter sets with a burn-in of 250,000 ignored initial parameter steps and a thin rate of ten ignored for every one recorded parameter set (see Section 1.1). Figure 4 shows a representative MH trajectory on the plane of parameters *k*_*r*2_ and *γ* for Model 2.

**Figure 4.**
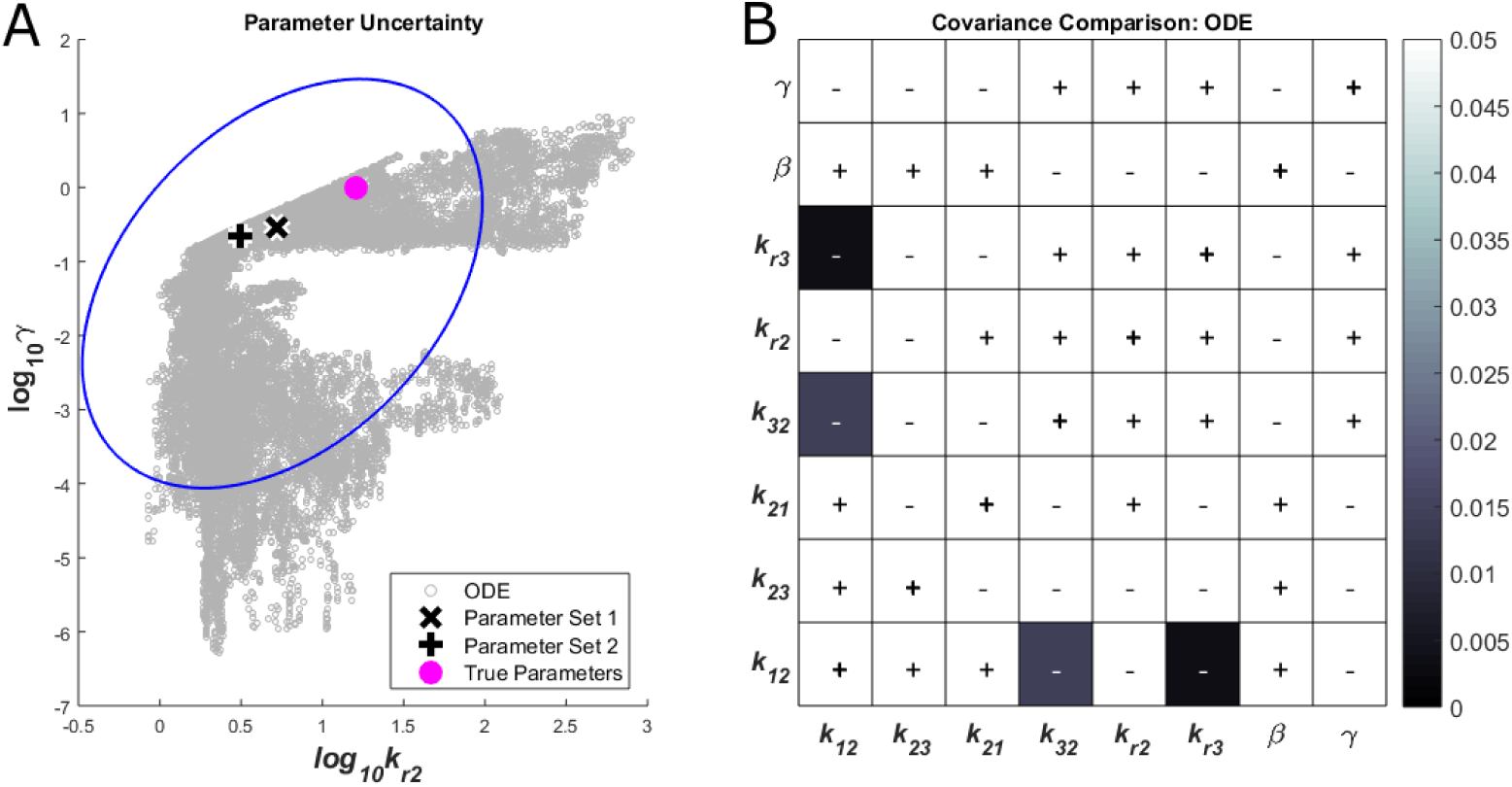
(A) Parameter uncertainty in logspace for parameters *k*_*r*2_ and *γ*, as determined by applying the Metropolis-Hastings algorithm to the ODE likelihood function (Eqn. 13). The ellipse indicates a 90% confidence interval, assuming a log Gaussian posterior. The magenta circle indicates the true *k*_*r*2_ and *γ* parameter values. Two similarly good parameter sets were selected from this chain: *log*_10_(**Λ**_**1**_) = [0.0705, −2.2859, −0.1497, −0.3444, 0.7181, −1.9591, −1.1805, −0.5304] and *log*_10_(**Λ**_**2**_) = [0.1797, 0.5551, −0.7058, −1.1098, 0.4949, 0.3303, 0.6639, −0.6531]. For these parameters sets, *k*_*r*2_ and *γ* are represented by the black **x** and **+**, respectively. (B) Comparison of the *thresholded* covariances for each parameter combination from the ODE MH search, with the covariance values thresholded at ±0.05. All covariance values whose magnitudes exceed ±0.05 are shown in white. The **(+)** and **(−)** symbols indicate positive and negative covariances, respectively.

Since the MH algorithm is random, each finite length run of this algorithm produces different results. In practice, the MH algorithm should be run until convergence (e.g., until multiple independent chains with different initial starting parameter guesses converge, within some specified metric and tolerance). Our analysis to compare the ODE model to the simulated data yielded the following average parameter values:

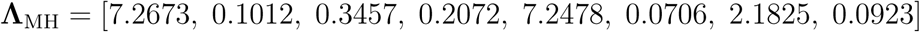

and the following large covariances for the parameters in logarithmic space:

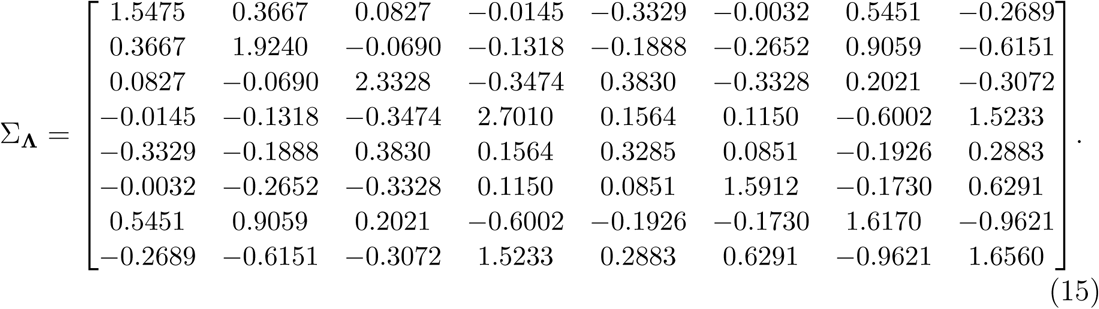

In Fig. 4, the ellipse depicts the 90% confidence interval in the parameters, with the (dubious) assumption that the posterior parameter distribution is a multivariate lognormal distribution with the mean equal to **Λ**_MH_ and a covariance matrix equal to Σ_**Λ**_. Clearly, with the data available, *the model is not well identified despite excellent fits to the average data*. This is an illustration of the effect of model sloppiness, where parameters are highly uncertain in some directions, yet potentially well-constrained in other directions within parameter space [39, 40]. In Section 4.5, we explore how stochastic analyses can dramatically improve model identification, even when using the same model and the same data.

### 4.4. Simulation of Stochastic Gene Expression Models

Next, we simulate many stochastic trajectories for the gene regulation model, thus generating potential single-cell mRNA distributions from the model. To do this, we define the stoichiometry matrix and the propensity functions for each of the four possible models. Unlike the standard SSA, one of the propensity functions in these models is a time-varying function. In this case, it is necessary to adapt the SSA in a manner similar to that discussed in [41, 42]. This modification corresponds to adding a fast reaction to the system, which has a large propensity function (*w*_1_ = 100*/*min), but which does not change the state of the system (**S**(:, 1) = **0**). We ran 1000 SSA trajectories for each of the four possible models, and we made histograms of the number of mRNA in different cells at each point in time (using the resulting best fit parameters from Section 4.2).

A single SSA trajectory is shown Fig. 5A. In Fig. 5B, we plot the average of multiple sets of 1000 SSA simulations versus time along with the ODE solution. In addition, Fig. 5C shows different FSP distributions for the two parameter sets (given in the caption for Fig. 4) and the simulated data at the same, specific time point. Despite producing excellent fits to the averaged data (Fig. 5B), neither parameter set captures the bimodal behavior of the data, as shown in Fig. 5C. This again shows that fitting the set of ODEs was not a sufficient strategy to extract all available information from our simulated single-cell data set, and the result suggests that fitting full distributions could lead to improvements in parameter identification [11]. We examine this possibility further in the next subsection.

**Figure 5.**
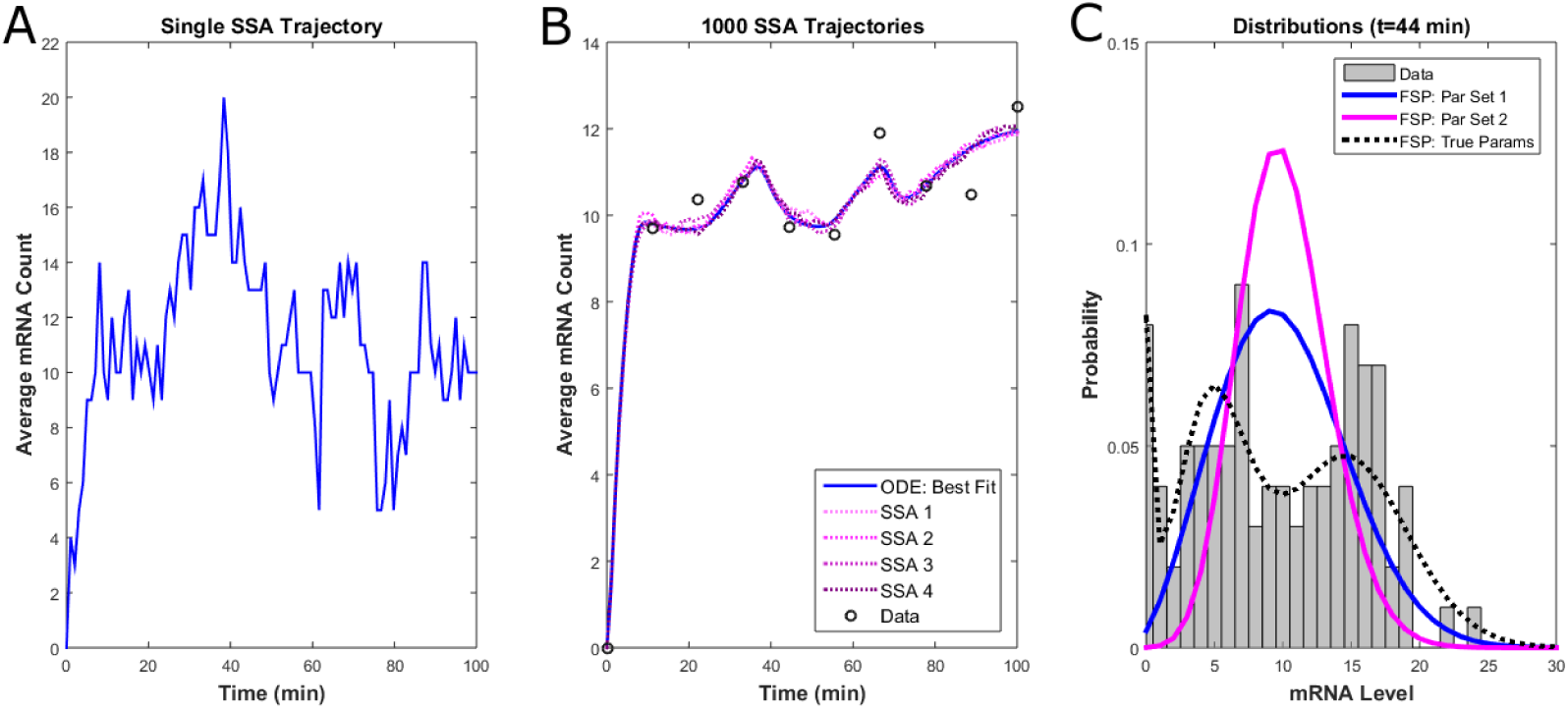
(A) Results from a single SSA trajectory for Model 2 using the best fit parameters, **Λ**_**Fit**_, obtained in Section 4.2. (B) Mean gene expression as a function of time from multiple runs of 1000 SSA trajectories using the best fit parameters compared to the deterministic solution and the experimental data. (C) Full distributions at 44 minutes for the two parameter sets, **Λ**_**1**_ and **Λ**_**2**_, compared to the data and the exact distribution for the true parameters at the same time point. The true parameters are **Λ**_**True**_ = [0.20, −0.076, 0.050, 0.050, 16.0, 5.0, 0.10, 1.0] or equivalently **log**_**10**_(**Λ**_**True**_) = [−0.6990, −1.1192, −1.3010, −1.3010, 1.2041, 0.6990, −1.0000, 0].

### 4.5. Parameter Uncertainty Estimation for FSP Models

To improve upon the parameter uncertainty estimation results, we next utilized the Finite State Projection (FSP) approach to re-explore the uncertainty in parameter space, given the single-cell data. To accomplish this, we write the CME in the matrix ODE form:

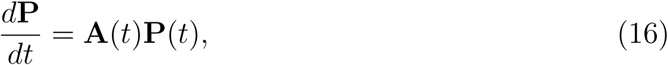

where **P** = [*p*_1_, *p*_2_, *…*] is the vector of probabilities of each state of the stochastic process, and **A**(*t*) is known as the *infinitesimal generator matrix*. See [14, 21, 22, 23] for examples on the construction and evaluation of this FSP formulation (Eqn. 16) and the GitHub folder at “https://github.com/MunskyGroup/WeberPB/Matlab_Codes” for specific FSP codes for this section. Equation 16 can then be integrated in time to compute full probability distributions for the gene regulation dynamics. We verified that the FSP analysis produced distributions that matched to the SSA results in the previous section (Fig. 5C, compare bars and dashed line). With the FSP-computed probability distributions, the likelihood of the data, given the model, is easily computed using the expression:

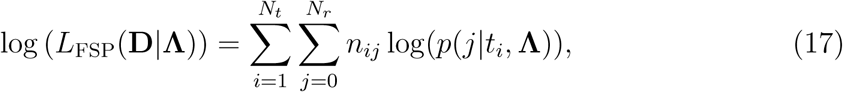

where *N*_*r*_ is the maximum number of mRNA observed in a single cell, *n*_*ij*_ is the number of cells with exactly *j* mRNA at the *i*^th^ time point, and *p*(*j|t*_*i*_, **Λ**) is the probability to have *j* mRNA at that time, given the model parameters in **Λ**, as discussed in [21, 23]. The corresponding log-likelihood of the model, given the data, can be written (using our previous definition of the prior) as:

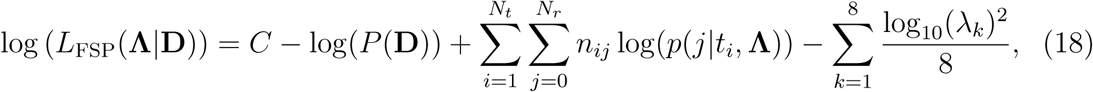

where *C* is a constant that is independent of the parameters. With this new likelihood function, we again implemented the MH algorithm to explore parameter uncertainty in the model, given the full information within the simulated single-cell data. In comparison to the parameter uncertainty estimation using the ODE analysis, our analysis using the FSP approach yielded substantially lower covariances for the parameters, as seen in the logarithmic covariance matrix:

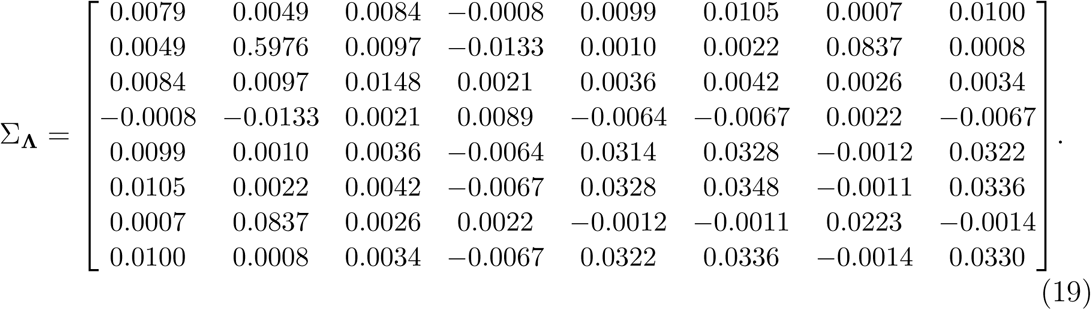

The FSP distribution results are presented in Figs. 6 and 7. Figure 6 shows that the FSP distributions resulting from the identified parameters match closely to the data distributions at the time points presented and capture the bimodal behavior of the data. Unlike the ODE analysis, the FSP was able to recover the correct model (i.e., Model 2) from its fit to the simulated data. Moreover, Fig. 7 shows that there is substantially less parameter uncertainty when the Metropolis-Hastings search is performed with the FSP instead of the ODE analysis. Figure 8 shows the true parameter values compared to the average estimated parameter values for Model 2 from the ODE and FSP searches of the parameter space. For nearly all of the parameters, the FSP search yields substantially closer estimates of the true parameter values and has much lower standard deviations for the parameter estimates as compared to ODE-based search. The only exception to this observation, for parameter *k*_23_(0), is easily explained by the fact that this reaction (Eqn. 2) is saturated and, therefore, unobservable for much of the system trajectory. It is also worth noting that, in addition to providing better parameter estimates and lower uncertainty, the FSP-based analysis required far shorter MH chains to explore parameter space before converging around the global minimum.

**Figure 6.**
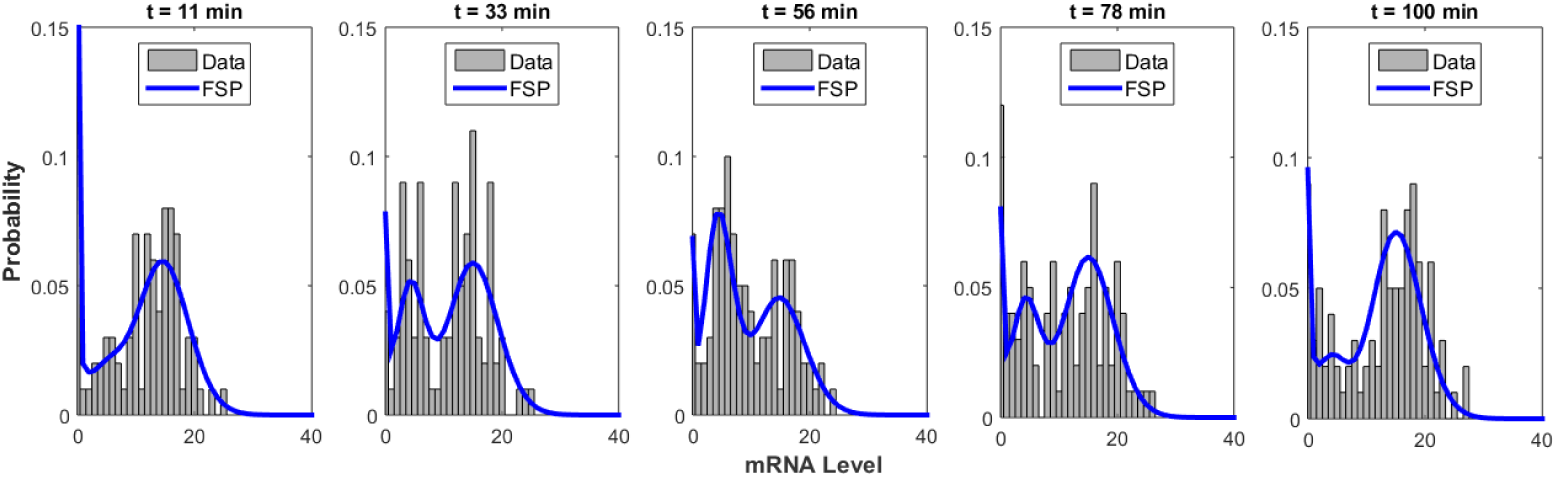
Data distributions compared with FSP distributions for Model 2 (using the FSP best fit parameter values) at specific time points.

**Figure 7.**
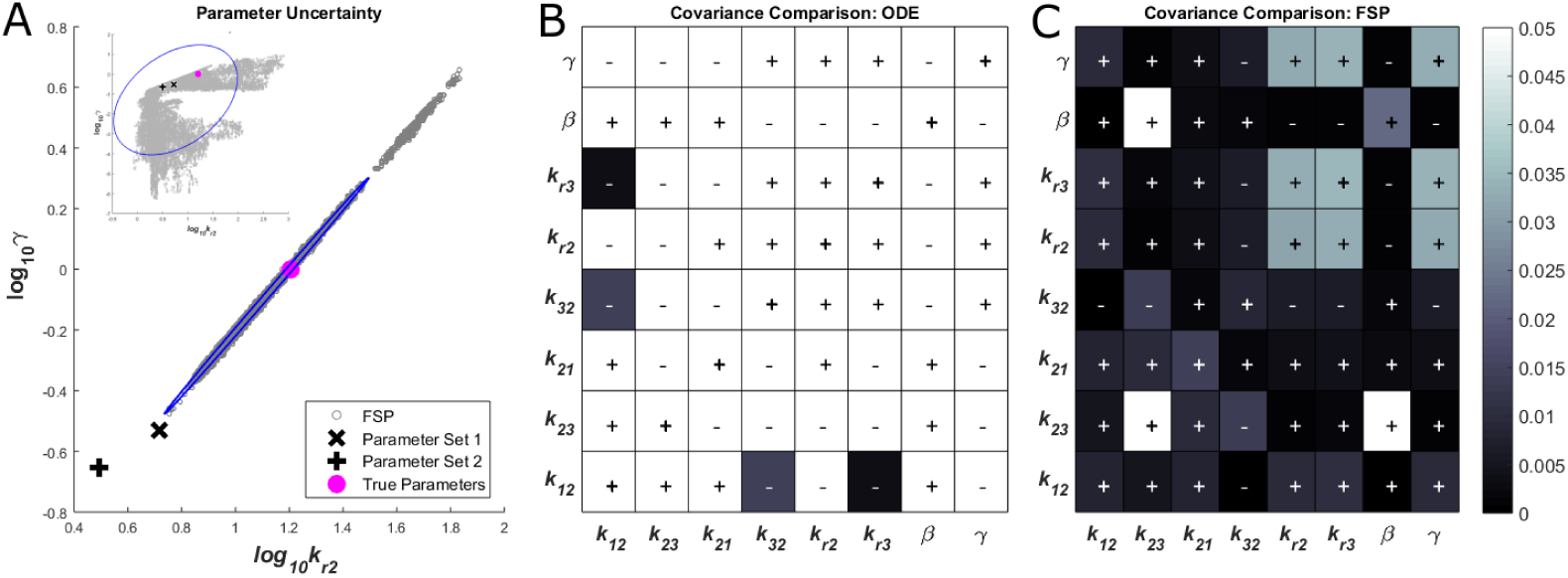
(A) Parameter uncertainty for *k*_*r*2_ and *γ*, as determined using the FSP likelihood function (Eqn. 18) in the Metropolis-Hastings analysis of Model 2. The ellipse indicates a 90% confidence interval. The magenta circle indicates the true *k*_*r*2_ and *γ* values. (A Inset) To provide a scale reference, the FSP-MH results are compared to the ODE search results for the same data set. (B,C) Comparison of the *thresholded* covariances for each parameter combination from the ODE MH search (B) and the FSP MH search (C), using the thresholding procedure described in the caption of Fig. 4 caption. The **(+)** indicates a positive covariance and **(−)** indicates a negative covariance.

**Figure 8.**
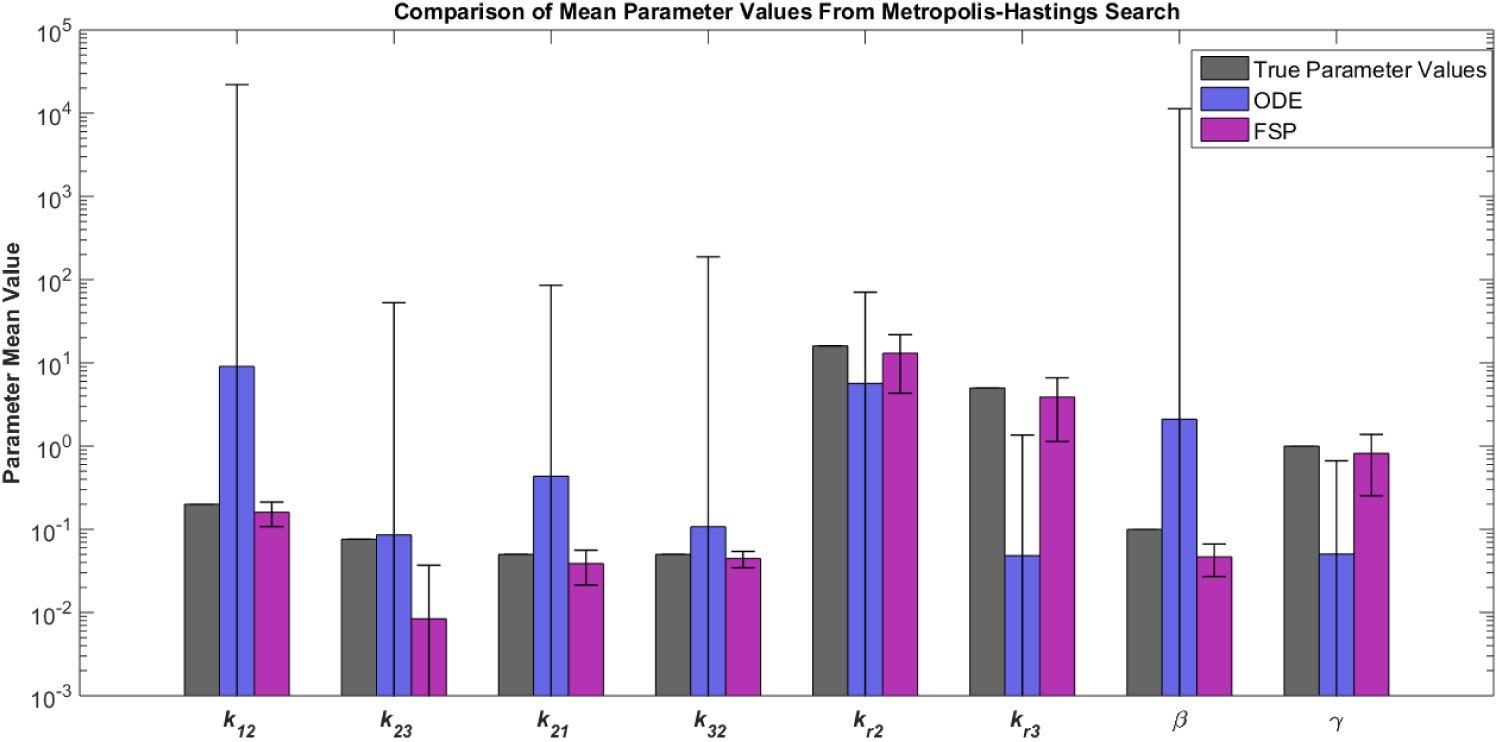
Bar plot showing the true parameter values compared to the mean parameter values from the ODE MH search and the FSP MH search for Model 2. The error bars represent the mean parameter value ± one standard deviation.

### 4.6. Directions of Parameter Uncertainty

To quantify and compare the model sloppiness after parameter estimation using the ODE and FSP analyses, we examined the eigenvectors and eigenvalues of the MCMC results (in logarithmic space). The square-root of the largest eigenvalue and its corresponding eigenvector quantify the magnitude and direction of the greatest component of parameter uncertainty (also known as the first principal component or PC). The second eigenvalue/eigenvector pair characterizes the next most variable direction, and so on. Figure 9A shows the magnitudes of the eight ranked PCs for the ODE (red) and FSP (black) analyses. The solid lines in Fig. 9A correspond to identification using the initial condition (*S*_1_ and *R* = 0) as presented above, and the dashed lines correspond to using a different initial condition defined by the steady-state distribution in the absence of external input. For this particular combination of input signal and sampling frequency, the two different initial conditions yield similar results for the contrast between the ODE and FSP analyses. However, we note that both the ODE and the FSP analyses are slightly more variable when the initial condition is defined by the steady-state behavior in the absence of the activating signal.

**Figure 9.**
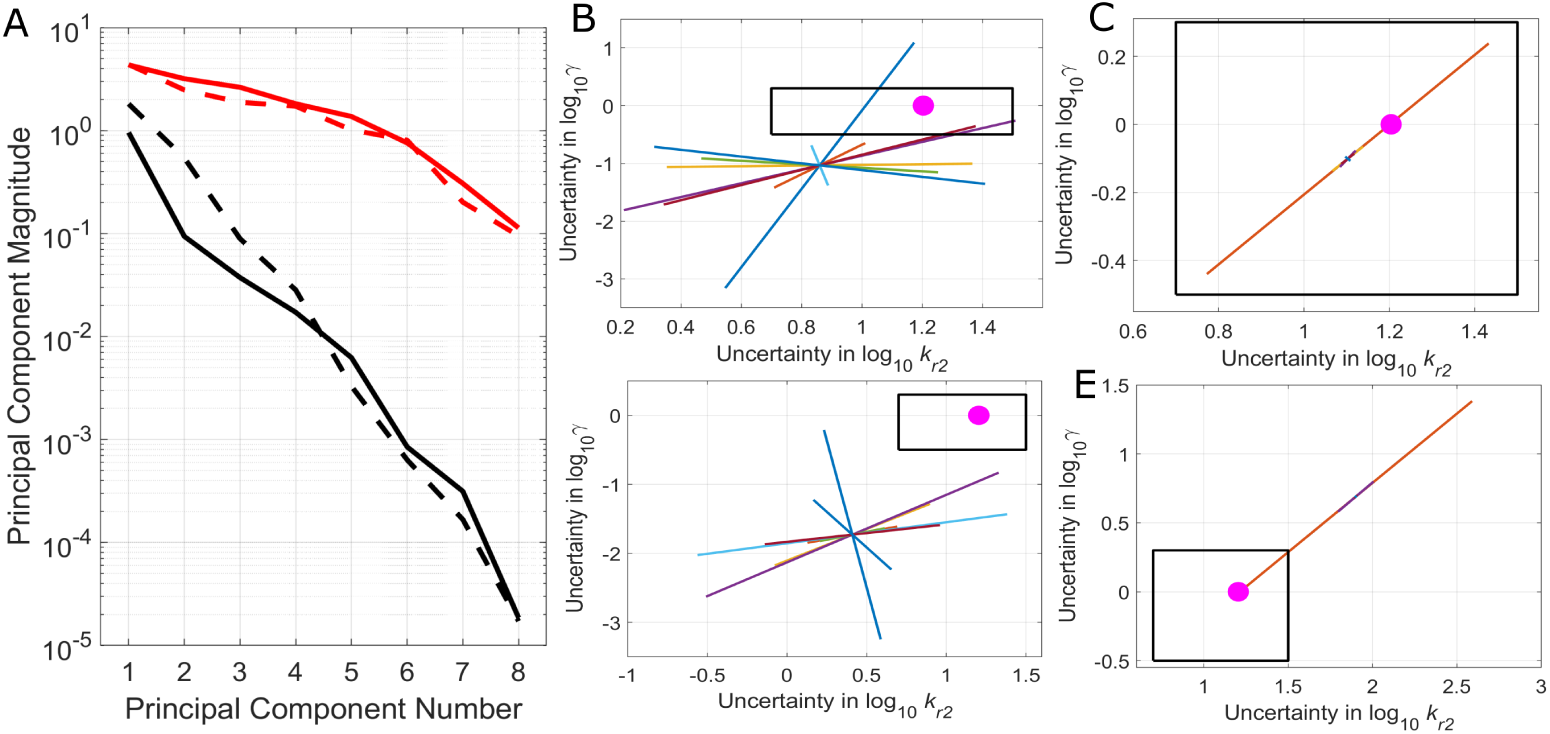
Principal component analysis of the posterior uncertainty after parameter estimation. (A) The magnitude of uncertainty (i.e., parameter variances) for the FSP (black) and ODE (red) analyses, ranked from the least to the best constrained directions in parameter space. Solid lines denote results using the initial conditions described in the text, and dashed lines denote the identification results assuming that the initial conditions are determined by the equilibrium condition in the absence of external signal. (B,C) Directions and magnitudes (*±* two standard deviations) of the principal components on the log10(*k*_*r*2_) versus log10(*γ*) plane using the (B) FSP analysis or (C) ODE analysis. Panels (B,C) use the initial conditions described in the text, and (D,E) use the zero-input steady state as the initial condition. For scale comparison of scales, the black box is the same size in every panel, and the magenta circle denotes the true parameter set.

Figures 9B-E show the directions of all eight PCs on the log_10_(*k*_*r*2_) *×* log_10_(*γ*) plane for the FSP and ODE analyses with the two different initial conditions. For the FSP analysis with the zero-activation initial condition (Fig. 9 B), the first PC explains 86% of the total uncertainty, and only three PCs are needed to explain more than 95% of the total uncertainty. Moreover, parameters *k*_*r*2_ and *γ* are linearly related to one another (i.e., their logarithms co-vary with a ratio of one) in each of the first three PCs, and Fig. 9B shows almost no uncertainty except on a single line with unit slope (Fig. 9C shows the same trend for the other initial condition). In contrast, six of the eight of the possible PCs are required in order to explain the total uncertainty in the ODE analysis, and Figs. 9C,E show that all of these PCs have significant variation in the log_10_(*k*_*r*2_) *×* log_10_(*γ*) plane, for either initial condition.

### 4.7. Importance of Experiment Design

Up to this point, we have demonstrated that the type of analysis chosen (i.e., an ODE- or FSP-based likelihood function) to examine experimental data is crucial in model and parameter identification. We have also observed that the type of initial condition can have a smaller, but still significant quantitative effect on parameter identification. We now briefly explore how different experimental designs can affect parameter identification results. To demonstrate this concern, we fit the data and performed MH searches of parameter space for multiple different experiments (i.e., different input signals). Specifically, we compared the original input signal, given in Eqn. 1, with two additional input signals, which are as follows:

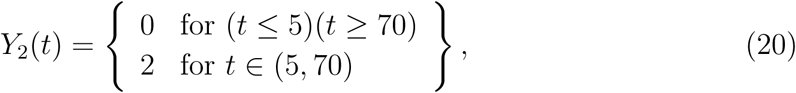

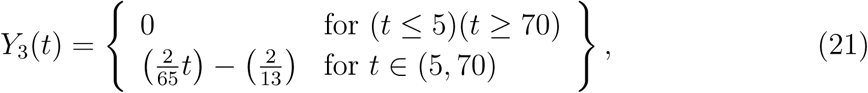

where both additional input signals begin at time *t* = 5 min and end at time *t* = 70 min, similar to *Y*_1_(*t*).

Figures 10 and 11 illustrate how the three different experimental designs (i.e., three different input signals) lead to dramatically different parameter uncertainties, using the ODE and the FSP likelihood functions, respectively. In all figures, we assume the exact same amount of data, but the effects on which parameters are tightly identified and which remain uncertain is non-trivially dependent on the input signal. Moreover, input signals that are best for one type of analysis may not be the best choice for a different analysis (for example, compare Figs. 10G to 11G).

**Figure 10.**
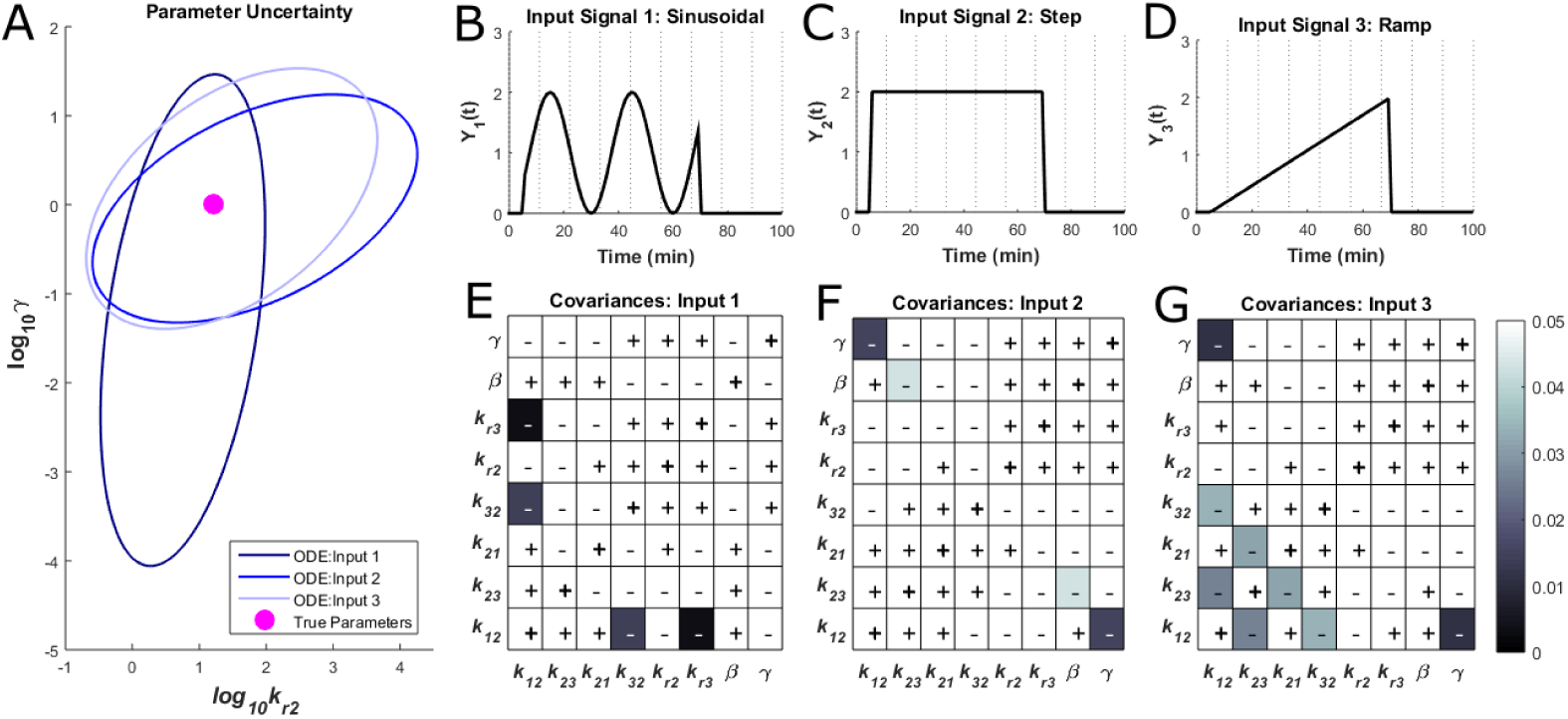
(A) Uncertainty in *k*_*r*2_ and *γ* given three different input signals, as determined by the ODE-based MH analysis of Model 2. The ellipses indicate a 90% confidence interval. The magenta circle indicates the true *k*_*r*2_ and *γ* parameter values. (B,C,D) Three different input signals and (E,F,G) the corresponding comparison of the *thresholded* covariances for each parameter combination from the ODE MH analysis. The **(+)** indicates positive covariances and **(−)** indicates negative covariances.

**Figure 11.**
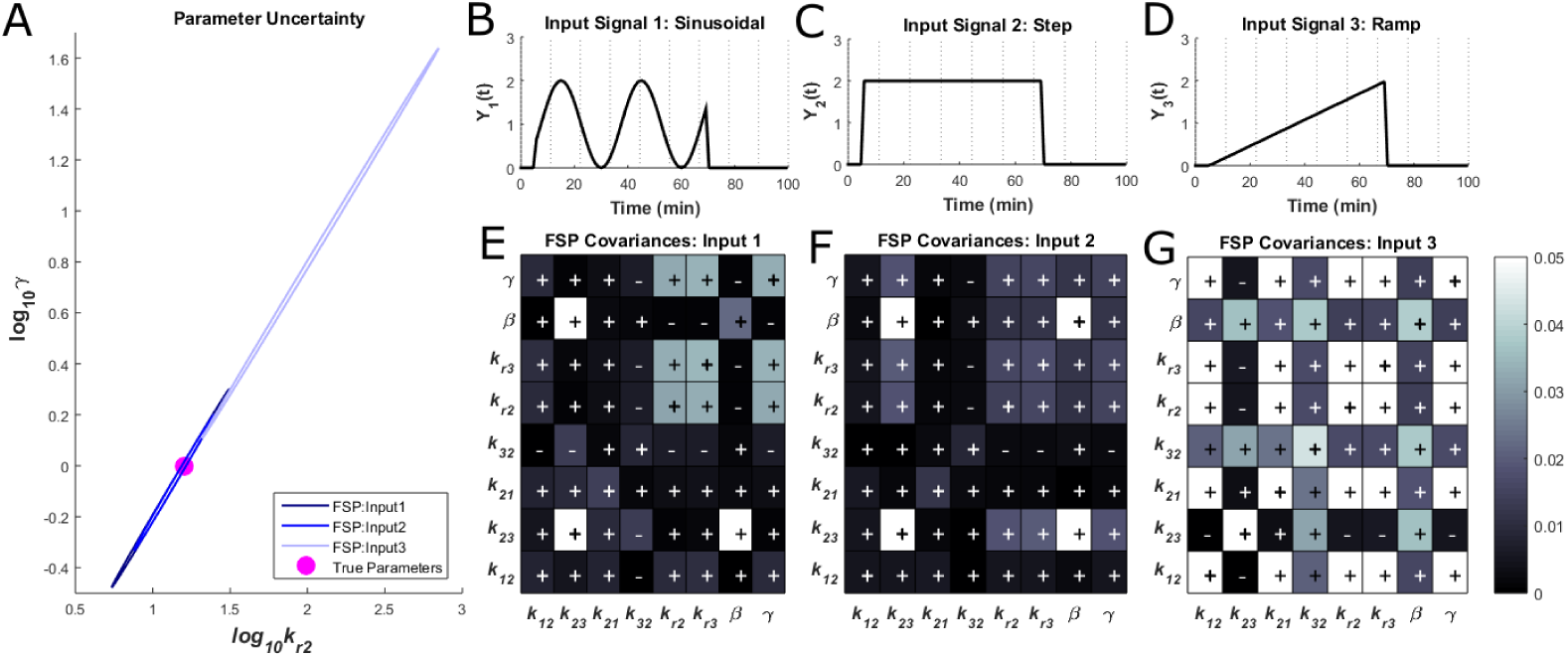
(A) Uncertainty in *k*_*r*2_ and *γ* given three different input signals, as determined by the FSP-based MH analysis of Model 2. The ellipses indicate a 90% confidence interval. The magenta circle indicates the true *k*_*r*2_ and *γ* parameter values. (B,C,D) Three different input signals and (E,F,G) the corresponding comparison of the *thresholded* covariances for each parameter combination from the FSP MH analysis. The **(+)** indicates positive covariances and **(−)** indicates negative covariances.

To further demonstrate how important experimental design considerations are to obtain accurate prediction of gene regulation dynamics, we make model predictions of FSP distributions at different times and for multiple different experiments (i.e., different input signals). In these predictions, we use the best-fit parameter sets from the FSP-based searches for all four models (each with a different time-dependent transition rate and obtained using the original input signal given in Eqn. 1). The potential input signals are *Y*_2_(*t*) (Eqn. 20), *Y*_3_(*t*) (Eqn. 21), and *Y*_4_(*t*), given by:

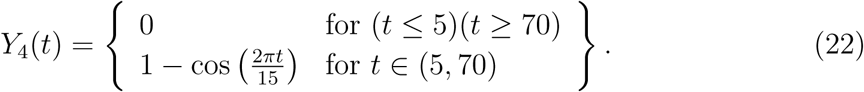

Figure 12 shows predictions for these 3 potential input signals (left, middle, right columns) for two potential time points (middle, bottom rows), assuming the 4 different models that were identified using the original experiment (different colors). By examining these predictions, we can gain insight into which experiments might be more informative. For example, at *t* = 30 min (middle row), an experiment with input signal *Y*_2_(*t*) (left column) or *Y*_3_(*t*) (middle column) would enable us to differentiate easily between the models, whereas input signal *Y*_4_(*t*) (right column) is unlikely to differentiate between the models. Furthermore, measuring the output at *t* = 30 min is clearly more informative than at *t* = 100 min (compare Figs. 12D-F with 12G-I).

**Figure 12.**
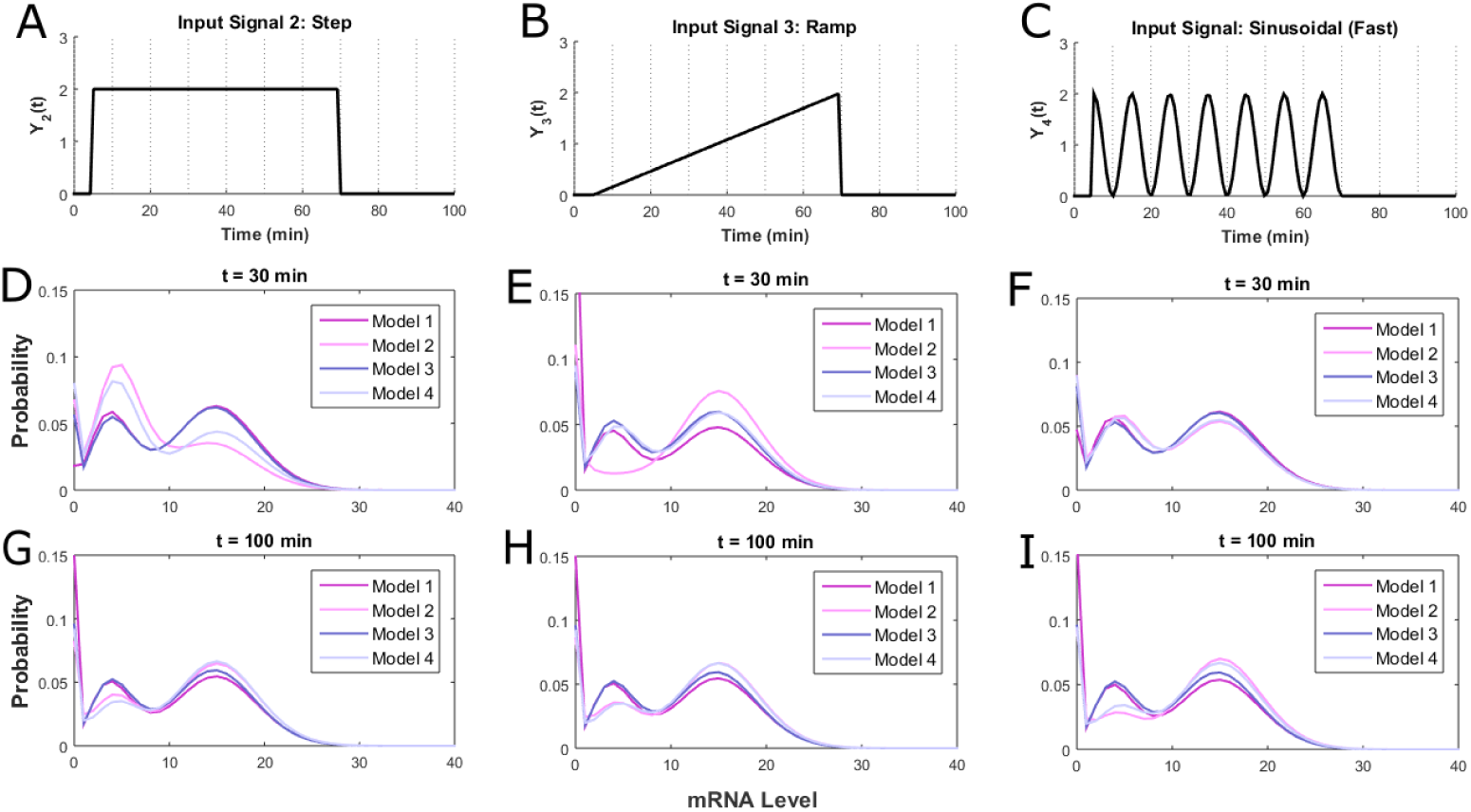
Predictions of the FSP distributions for all four model numbers at two different time points for three different experiments (with input signals shown in Eqn. 20 - 22). Each model uses the best parameter set found during the fit to the original input signal in Eqn. 1.

## 5. Summary and Conclusions

In this article, we explored the application of various computational techniques (such as ODE analyses [43, 44, 45, 46, 47, 48], stochastic simulations [17, 49, 50, 51, 52, 53, 54, 55, 56, 57, 58, 59, 60], and finite state projections [12, 14, 20, 21, 23, 61, 62, 18, 63, 64]) to understand the dynamics of single-cell gene regulation. In particular, we focused on a class of simplified multi-state gene regulation models that has previously been fit to experimental single-cell data [1, 10, 11], and we used a combination of simulated single-cell data and different modeling procedures to explore how well one could identify models and parameter uncertainties from discrete stochastic data. Additionally, we discussed the importance of experiment design and illustrated how different initial conditions or experiments can be more informative than others for model identification. In addition to changing the initial conditions or input conditions, additional information can be gained from system response by adjusting the times at which measurements are taken. Although there have been many recent developments toward model-driven experiment design for ODE analyses [25, 65], the development of experiment design methods for stochastic gene regulatory systems remains an active field of investigation [25, 66]. Moreover, our analyses show that fitting ODEs to dynamical data may not be adequate for parameter identification and discerning parameter uncertainty (even when models are relatively simple), and we discussed the fact that fitting full distributions (e.g., using an FSP likelihood function) may lead to substantial improvements, even when considering the same model and same data. These results argue that the integration of discrete stochastic models and single-cell data could open new possibilities for improved model identification, even in cases where traditional ODE analyses have failed due to excessive parameter sloppiness.

In our current study, we focused on the analysis of mRNA levels as can be measured precisely with single-molecule resolution and fast temporal resolution using single-molecule Fluorescence *in situ* Hybridization [1, 5, 9, 11]. Similar tools and stochastic model can also be applied to the analysis of protein variations, although such analyses must contend with additional complications, such as slow protein translation or activation dynamics that can filter out the fast transcriptional responses [13] or added background noise (e.g., cellular autofluorescence) that must be convolved with the model-generated protein signal [67]. Despite these issues, single-cell flow cytometry data have been successfully combined with stochastic models to generate accurate predictive models for single-cell protein distributions [67, 68]. We assumed the possibility of rich and well measured time-varying input signals, which can be temporally controlled through alteration of environmental conditions and quantified with time-lapse fluorescence microscopy [1, 10, 31, 32, 33].

To make these approaches more accessible, we have developed a GUI, which is freely available on GitHub in the folder “https://github.com/MunskyGroup/WeberPB/GUI”. The GUI, shown in Fig. 13, provides an opportunity for users to specify any input and model number and to perform the analyses outlined throughout, evaluate results, and generate figures. In addition, the GUI enables the automatic generation of simulated data using SSA for user-specified parameter sets and models, or to import a previously generated data file to perform the analyses outlined in this exercise. The GUI requires Python and the packages: NumPy, SciPy, and Matplotlib.

**Figure 13.**
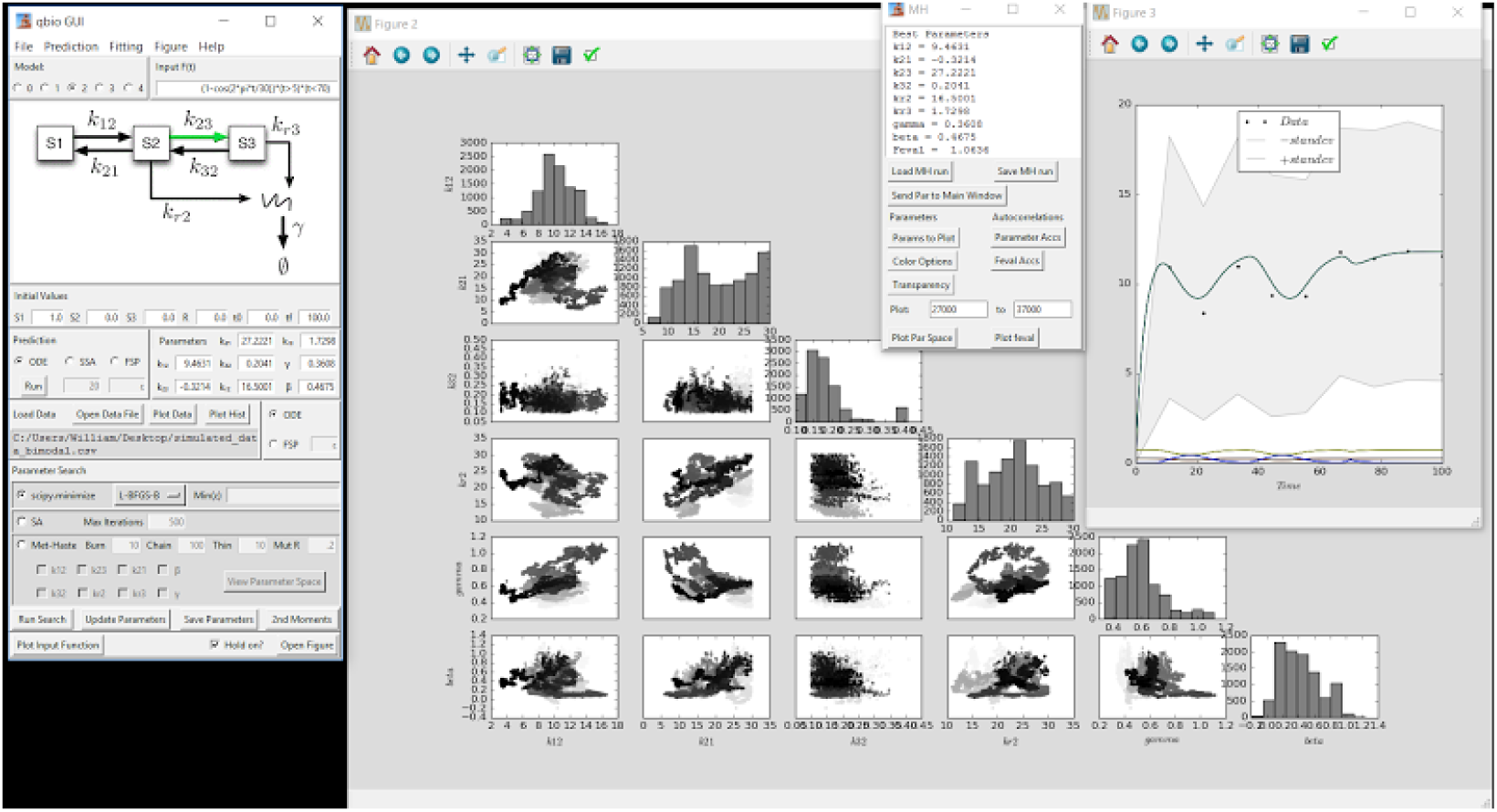
Screenshot of the provided GUI, which can be utilized to perform all analyses outlined throughout, including generating simulated data sets and producing desired plots.

## Acknowledgements

BM and LW were funded by the W.M. Keck Foundation. BM and WR were funded by NIH R35GM124747.

## Footnotes

‡ A substantially reduced version of this manuscript will appear as an extended exercise in the textbook *Quantitative Biology: Theory, Computational Methods and Examples of Models*, edited by Brian Munsky, Lev S. Tsimring, and William S. Hlavacek, to be published by MIT Press, [3].

## References

[1] G. Neuert, B. Munsky, R. Z. Tan, L. Teytelman, M. Khammash, and A. van Oudenaarden, “Systematic Identification of Signal-Activated Stochastic Gene Regulation,” Science, vol. 339, pp. 584–587, Jan. 2013.

[2] A. Senecal, B. Munsky, F. Proux, N. Ly, F. E. Braye, C. Zimmer, F. Mueller, and X. Darzacq, “Transcription factors modulate c-fos transcriptional bursts,” Cell reports, vol. 8, no. 1, pp. 75–83, 2014.

[3] B. Munsky, W. S. Hlavacek, and L. S. Tsimring, Quantitative Biology: Theory, Computational Methods and Examples of Models. MIT Press, 1st ed., In Press.

[4] T. Morisaki, K. Lyon, K. F. DeLuca, J. G. DeLuca, B. P. English, Z. Zhang, L. D. Lavis, J. B. Grimm, S. Viswanathan, L. L. Looger, et al., “Real-time quantification of single rna translation dynamics in living cells,” Science, vol. 352, no. 6292, pp. 1425–1429, 2016.

[5] A. Raj, P. van den Bogaard, S. A. Rifkin, A. van Oudenaarden, and S. Tyagi, “Imaging individual mRNA molecules using multiple singly labeled probes,” Nature Methods, vol. 5, pp. 877–879, Sept. 2008.

[6] H. M. Shapiro, Practical flow cytometry. John Wiley & Sons, 2005.

[7] M. H. Spitzer and G. P. Nolan, “Mass cytometry: single cells, many features,” Cell, vol. 165, no. 4, pp. 780–791, 2016.

[8] C. Gawad, W. Koh, and S. R. Quake, “Single-cell genome sequencing: current state of the science,” Nature Reviews Genetics, vol. 17, no. 3, p. 175, 2016.

[9] A. M. Femino, F. S. Fay, K. Fogarty, and R. H. Singer, “Visualization of Single RNA Transcripts in Situ,” Science, vol. 280, pp. 585–590, Apr. 1998.

[10] A. Senecal, B. Munsky, F. Proux, N. Ly, F. E. Braye, C. Zimmer, F. Mueller, and X. Darzacq, “Transcription factors modulate c-Fos transcriptional bursts.,” Cell Reports, vol. 8, pp. 75–83, July 2014.

[11] B. Munsky, G. Li, Z. Fox, D. P. Shepherd, and G. Neuert, “Distribution shapes govern the discovery of predictive models for gene regulation,” bioRxiv, p. 154401, 2017.

[12] B. Munsky, G. Neuert, and A. van Oudenaarden, “Using Gene Expression Noise to Understand Gene Regulation,” Science, vol. 336, pp. 183–187, Apr. 2012.

[13] B. Munsky and G. Neuert, “From analog to digital models of gene regulation,” Physical Biology, vol. 12, no. 4, p. 045004, 2015.

[14] B. Munsky, Z. Fox, and G. Neuert, “Integrating single-molecule experiments and discrete stochastic models to understand heterogeneous gene transcription dynamics,” Methods, vol. 85, pp. 12–21, Sept. 2015.

[15] S. Chib and E. Greenberg, “Understanding the Metropolis-Hastings Algorithm,” The American Statistician, vol. 49, pp. 327–335, Nov. 1995.

[16] D. T. Gillespie, “A general method for numerically simulating the stochastic time evolution of coupled chemical reactions,” Journal of Computational Physics, vol. 22, no. 4, pp. 403–434, 1976.

[17] D. T. Gillespie, “Exact stochastic simulation of coupled chemical reactions,” The journal of physical chemistry, vol. 81, no. 25, pp. 2340–2361, 1977.

[18] D. A. McQuarrie, “Stochastic approach to chemical kinetics,” Journal of Applied Probability, vol. 4, no. 3, pp. 413–478, 1967.

[19] N. van Kampen, Stochastic Processes in Physics and Chemistry. Elsevier, 3 ed., 2007.

[20] B. Munsky and M. Khammash, “The finite state projection algorithm for the solution of the chemical master equation,” J. Chem. Phys., vol. 124, no. 4, p. 044104, 2006.

[21] Z. Fox, G. Neuert, and B. Munsky, “Finite state projection based bounds to compare chemical master equation models using single-cell data,” The Journal of chemical physics, vol. 145, p. 074101, Aug. 2016.

[22] K. N. Dinh and R. B. Sidje, “Understanding the finite state projection and related methods for solving the chemical master equation,” Physical biology, vol. 13, no. 3, p. 035003, 2016.

[23] Z. Fox and B. Munsky, “Stochasticity or noise in biochemical reactions,” in Quantitative Biology: Theory, Computational Methods and Examples of Models (B. Munsky, L. S. Tsimring, and W. S. Hlavacek, eds.), ch. 5, Cambridge, Massachusetts: MIT Press, 2018.

[24] K. NienaĆtowski, T. Jetka, and M. Komorowski, “Sensitivity analysis,” in Quantitative Biology: Theory, Computational Methods and Examples of Models (B. Munsky, L. S. Tsimring, and W. S. Hlavacek, eds.), Cambridge, Massachusetts: MIT Press, 2018.

[25] T. Jetka, K. NienaCtowski, and M. Komorowski, “Experimental design,” in Quantitative Biology: Theory, Computational Methods and Examples of Models (B. Munsky, L. S. Tsimring, and W. S. Hlavacek, eds.), Cambridge, Massachusetts: MIT Press, 2018.

[26] B. C. Daniels, M. Dobrzynski, and D. Fey, “Parameter estimation, sloppiness, and model identifiability,” in Quantitative Biology: Theory, Computational Methods and Examples of Models (B. Munsky, L. S. Tsimring, and W. S. Hlavacek, eds.), Cambridge, Massachusetts: MIT Press, 2018.

[27] J. Bernie J. Daigle, “Bayesian parameter estimation and markov chain monte carlo,” in Quantitative Biology: Theory, Computational Methods and Examples of Models (B. Munsky, L. S. Tsimring, and W. S. Hlavacek, eds.), Cambridge, Massachusetts: MIT Press, 2018.

[28] B. Mélykúti, E. August, A. Papachristodoulou, and H. El-Samad, “Discriminating between rival biochemical network models: three approaches to optimal experiment design,” BMC Systems Biology, vol. 4, no. 1, p. 1, 2010.

[29] J. F. Apgar, J. E. Toettcher, D. Endy, F. M. White, and B. Tidor, “Stimulus Design for Model Selection and Validation in Cell Signaling,” PLoS Computational Biology, vol. 4, p. e30, Feb. 2008.

[30] J. Peccoud and B. Ycart, “Markovian Modeling of Gene-Product Synthesis,” Theoretical Population Biology, vol. 48, pp. 222–234, Oct. 1995.

[31] D. Muzzey, C. A. Gómez-Uribe, J. T. Mettetal, and A. van Oudenaarden, “A Systems-Level Analysis of Perfect Adaptation in Yeast Osmoregulation,” Cell, vol. 138, pp. 160–171, July 2009.

[32] J. T. Mettetal, D. Muzzey, C. Gomez-Uribe, and A. van Oudenaarden, “The Frequency Dependence of Osmo-Adaptation in Saccharomyces cerevisiae,” Science, vol. 319, pp. 482–484, Jan. 2008.

[33] G. Neuert, “Dynamic control and model inference of signal activated gene regulation,” in APS March Meeting Abstracts, 2017.

[34] B. Munsky and M. Khammash, “The Finite State Projection Algorithm for the Solution of the Chemical Master Equation,” The Journal of Chemical Physics, vol. 124, no. 4, p. 044104, 2006.

[35] M. Krzywinski and N. Altman, “Points of significance: importance of being uncertain,” 2013.

[36] J. Nelder and R. Mead, “A Simplex Method for Function Minimization,” The Computer Journal, vol. 7, no. 4, p. 308, 1965.

[37] G. B. Dantzig, Linear Programming and Extensions. Princeton, NJ: Princeton University Press, 1963.

[38] P. J. Van Laarhoven and E. H. Aarts, “Simulated annealing,” in Simulated annealing: Theory and applications, pp. 7–15, Springer, 1987.

[39] R. N. Gutenkunst, J. J. Waterfall, F. P. Casey, K. S. Brown, C. R. Myers, and J. P. Sethna, “Universally sloppy parameter sensitivities in systems biology models.,” PLoS computational biology, vol. 3, pp. 1871–78, Oct. 2007.

[40] B. C. Daniels, Y.-J. Chen, J. P. Sethna, R. N. Gutenkunst, and C. R. Myers, “Sloppiness, robustness, and evolvability in systems biology.,” Current Opinion in Biotechnology, vol. 19, no. 4, pp. 389–395, 2008.

[41] M. Voliotis, P. Thomas, C. G. Bowsher, and R. Grima, “The extra reaction algorithm for stochastic simulation of biochemical reaction systems in fluctuating environments,” in Quantitative Biology: Theory, Computational Methods and Examples of Models (B. Munsky, L. S. Tsimring, and W. S. Hlavacek, eds.), Cambridge, Massachusetts: MIT Press, 2018.

[42] M. Voliotis, P. Thomas, R. Grima, and C. G. Bowsher, “Stochastic simulation of biomolecular networks in dynamic environments,” PLoS computational biology, vol. 12, no. 6, p. e1004923, 2016.

[43] D. Fey, M. Dobrzynski, and B. N. Kholodenko, “Modeling with ordinary differential equations,” in Quantitative Biology: Theory, Computational Methods and Examples of Models (B. Munsky, L. S. Tsimring, and W. S. Hlavacek, eds.), Cambridge, Massachusetts: MIT Press, 2018.

[44] E. D. Conrad and J. J. Tyson, System Modeling in Cellular Biology: From Concepts to Nuts and Bolts, ch. Modeling Molecular Interaction Networks with Nonlinear Ordinary Differential Equations, p. 97ff. The MIT Press, 2006.

[45] B. N. Kholodenko, O. V. Demin, G. Moehren, and J. B. Hoek, “Quantification of short term signaling by the epidermal growth factor receptor.,” J Biol Chem, vol. 274, pp. 30169–30181, Oct 1999.

[46] E. Klipp, W. Liebermeister, C. Wierling, A. Kowald, H. Lehrach, and R. Herwig, Systems Biology. Wiley, 2013.

[47] B. Ingalls, Mathematical Modeling in Systems Biology: An Introduction. MIT Press, 2013.

[48] J. Keener and J. Sneyd, Mathematical Physiology, vol. 8 of Interdisciplinary Applied Mathematics. New York: Springer-Verlag, second ed., 2001.

[49] R. Bertolusso and M. Kimmel, “Kinetic monte carlo analyses of discrete biomolecular events,” in Quantitative Biology: Theory, Computational Methods and Examples of Models (B. Munsky, L. S. Tsimring, and W. S. Hlavacek, eds.), Cambridge, Massachusetts: MIT Press, 2018.

[50] M. A. Gibson and J. Bruck, “Efficient exact stochastic simulation of chemical systems with many species and many channels,” Journal of Physical Chemistry A, vol. 104, pp. 1876–1889, 2000.

[51] A. Slepoy, A. P. Thompson, and S. J. Plimpton, “A constant-time kinetic Monte Carlo algorithm for simulation of large biochemical reaction networks.,” J Chem Phys, vol. 128, p. 205101, May 2008.

[52] D. Gillespie, “Approximate accelerated stochastic simulation of chemically reacting systems,” The Journal of Chemical Physics, vol. 115, no. 4, pp. 1716–1733, 2001.

[53] D. Gillespie, “The chemical Langevin equation,” The Journal of Chemical Physics, vol. 113, no. 1, pp. 297–306, 2000.

[54] D. Gillespie and L. Petzold, “Improved leap-size selection for accelerated stochastic simulation,” The Journal of Chemical Physics, vol. 119, no. 16, pp. 8229–8234, 2003.

[55] Y. Cao, D. Gillespie, and L. Petzold, “Efficient step size selection for the tau-leaping simulation method,” The Journal of Chemical Physics, vol. 124, no. 4, p. 044109, 2006.

[56] Y. Cao, D. Gillespie, and L. Petzold, “Adaptive explicit-implicit tau-leaping method with automatic tau selection,” The Journal of Chemical Physics, vol. 126, no. 22, p. 224101, 2007.

[57] Y. Cao, D. Gillespie, and L. Petzold, “The Slow-Scale Stochastic Simulation Algorithm,” The Journal of chemical physics, vol. 122, Jan. 2005.

[58] C. Rao and A. P. Arkin, “Stochastic Chemical Kinetics and the Quasi-Steady-State Assumption: Application to the Gillespie Algorithm,” The Journal of chemical physics, vol. 118, pp. 4999–5010, 2003.

[59] L. Harris and P. Clancy, “A “partitioned leaping” approach for multiscale modeling of chemical reaction dynamics,” J. Chem. Phys., vol. 125, p. 144107, 2006.

[60] Y. Cao, D. T. Gillespie, and L. R. Petzold, “Efficient step size selection for the tau-leaping simulation method,” The Journal of chemical physics, vol. 124, no. 4, p. 044109, 2006.

[61] K. Burrage, M. Hegland, S. Macnamara, and R. Sidje, “A Krylov-Based Finite State Projection Algorithm for Solving the Chemical Master Equation Arising in the Discrete Modelling of Biological Systems,” Proc. of The A.A.Markov 150th Anniversary Meeting, pp. 21–37, 2006.

[62] B. Munsky and M. Khammash, “The Finite State Projection Approach for the Analysis of Stochastic Noise in Gene Networks,” IEEE Trans. Automat. Contr./IEEE Trans. Circuits and Systems: Part 1, vol. 52, pp. 201–214, Jan. 2008.

[63] B. Munsky and M. Khammash, “Transient Analysis of Stochastic Switches and Trajectories with Applications to Gene Regulatory Networks,” IET Systems Biology, vol. 2, no. 5, pp. 323–333, 2008.

[64] B. Munsky, “Modeling cellular variability,” Quantitative Biology: From Molecular to Cellular Systems, p. 233, 2012.

[65] J. Liepe, S. Filippi, M. Komorowski, and M. P. Stumpf, “Maximizing the information content of experiments in systems biology,” PLoS Comput Biol, vol. 9, no. 1, p. e1002888, 2013.

[66] J. Ruess, A. Milias-Argeitis, and J. Lygeros, “Designing experiments to understand the variability in biochemical reaction networks,” Journal of The Royal Society Interface, vol. 10, no. 88, p. 20130588, 2013.

[67] B. Munsky, B. Trinh, and M. Khammash, “Listening to the noise: random fluctuations reveal gene network parameters.,” Molecular Systems Biology, vol. 5, no. 318, p. 318, 2009.

[68] C. Lou, B. Stanton, Y.-J. Chen, B. Munsky, and C. A. Voigt, “Ribozyme-based insulator parts buffer synthetic circuits from genetic context.,” Nature Biotechnology, vol. 30, pp. 1137–1142, Nov. 2012.

